# Electrophysiological correlates of attention in the locus coeruleus - anterior cingulate cortex circuit during the rodent continuous performance test

**DOI:** 10.1101/2023.04.19.537406

**Authors:** Henry L. Hallock, Suhaas Adiraju, Jorge Miranda-Barrientos, Jessica M. McInerney, Seyun Oh, Adrienne C. DeBrosse, Ye Li, Gregory V. Carr, Keri Martinowich

## Abstract

Sustained attention, the ability to focus on an activity or stimulus over time, is significantly impaired in many psychiatric disorders, and there remains a major unmet need in treating impaired attention. Continuous performance tests (CPTs) were developed to measure sustained attention in humans, non-human primates, rats, and mice, and similar neural circuits are engaged across species during CPT performance, supporting their use in translational studies to identify novel therapeutics. Here, we identified electrophysiological correlates of attentional performance in a touchscreen-based rodent CPT (rCPT) in the locus coeruleus (LC) and anterior cingulate cortex (ACC), two inter-connected regions that are implicated in attentional processes. We used viral labeling and molecular techniques to demonstrate that neural activity is recruited in LC-ACC projections during the rCPT, and that this recruitment increases with cognitive demand. We implanted male mice with depth electrodes within the LC and ACC for local field potential (LFP) recordings during rCPT training, and identified an increase in ACC delta and theta power, and an increase in LC delta power during correct responses in the rCPT. We also found that the LC leads the ACC in theta frequencies during correct responses while the ACC leads the LC in gamma frequencies during incorrect responses. These findings may represent translational biomarkers that can be used to screen novel therapeutics for drug discovery in attention.

## INTRODUCTION

Sustained attention, the ability to focus on a stimulus for long periods of time, is frequently disrupted in patients diagnosed with, as well as those at high genetic risk for a range of neuropsychiatric disorders. This includes children [1,2], and adults [3] with attention-deficit hyperactivity disorder (ADHD), patients in remission from major depressive disorder [4,5], as well as adults diagnosed with schizophrenia [6–8] and children at risk for schizophrenia [9]. Sustained attention in individuals with these disorders is commonly measured with continuous performance tests (CPTs) [10]. To successfully perform CPTs, subjects must focus their attention on an auditory [11] or visual stimulus. If one stimulus type is presented (the “target”), the subject must initiate a “go” response (for example, pressing a lever [10]). If a second stimulus type is presented (the “non-target”), subjects must withhold responding. Subjects can therefore make errors of omission (failing to respond to a target), or errors of commission (responding to a non-target), which may differentiate underlying deficits in motivation or cognitive control [12,13]. CPTs are highly effective tools for assessing sustained attention deficits in patient populations, as CPT performance is a much better predictor of clinical ADHD diagnosis than parent reports of ADHD symptoms [14]. Despite widespread use of CPTs as sustained attention assays, the neural mechanisms that govern performance are not well-understood.

Investigating the neural correlates of behavior during CPTs has been impeded by several barriers. First, multiple versions of CPTs exist [15–19], which complicates attempts to identify brain activity patterns that might generalize across experimental settings. Second, it is inherently difficult to obtain brain activity data in humans that is both spatially and temporally resolved without using invasive techniques. To overcome these barriers, a standardized touchscreen-based rodent version of the CPT (rCPT) was developed using task parameters that influence sustained attention in human CPTs [20–22]. The rCPT provides the opportunity to study how brain function contributes to sustained attention in a task with high translational potential; indeed, rodent models of developmental disruption with schizophrenia-relevant phenotypes are reliably impaired in the rCPT [23,24]. Identifying biomarkers in this task would therefore be invaluable for advancing diagnostic criteria and novel treatment strategies for the myriad disorders that feature sustained attention deficits.

The anterior cingulate cortex (ACC) is active during sustained attention tasks in humans, particularly under conditions of high attentional load (i.e., when distractors are present [25–28]). In concert with these findings, ACC activity is diminished during CPT performance in individuals diagnosed with ADHD [3,29], and schizophrenia [30,31], suggesting that the ACC is a neuroanatomical locus for attentional processing. Recent studies suggest that the primate ACC may be most anatomically and functionally homologous with the area commonly referred to as the medial prefrontal cortex (mPFC) in the rodent [32,33]. This is important because these anatomical discrepancies create obstacles to translating neural data between species [34]. The prelimbic (PrL) cortex, which is included in what is commonly termed mPFC, shows functional and anatomical similarity to the human dorsal ACC, which plays critical roles in attentional processes [28,35]. In the rodent, neurons in the ACC increase their firing rates prior to correct trials in sustained attention tasks [36–38], indicating that the ACC’s role in sustained attention is phylogenetically conserved across species. The rodent ACC is also implicated in rCPT performance [37,38], but the neurobiological mechanisms by which this brain region controls behavior in this task is, to our knowledge, unexplored.

To assess the relationship between ACC function and rCPT performance with circuit-level detail, we investigated recruitment of neural activity in projections to the ACC from the locus coeruleus (LC), a brain region that is heavily implicated in attention. We found that neurons in the LC with direct axonal input to the ACC are recruited during the rCPT, and that this recruitment increases with cognitive demand. To further query activity in the LC-ACC circuit during rCPT performance, we simultaneously recorded local field potentials (LFPs) in the LC and ACC and found an increase in ACC delta and theta power, as well as LC delta power following correct responses in well-trained mice. Moreover, LC neuronal activity leads ACC activity during correct responses in delta frequencies while ACC activity leads the LC during incorrect responses in gamma frequencies.

## MATERIALS AND METHODS

Abbreviated methods are provided here - full details provided in the supplement.

### Animals

8-10 week old male C57BL/6J mice (Strain#: 000664, The Jackson Laboratory) were group-housed (4/cage) until electrode implantation and then double-housed. Procedures were approved by the Johns Hopkins Animal Care and Use Committee.

### Food restriction protocol

Food was restricted to 2.5 g chow/mouse/day, and mice were maintained at 85-90% of predicted free-feeding weight.

### Surgical procedures

For immunohistochemistry experiments, holes were drilled with a 0.9 mm burr above the ACC (A/P:±1.7mm, M/L:+0.3mm), and a retrograde virus expressing tdTomato (AAVrg-CAG-tdTomato, Addgene catalog # 59462-AAVrg) was injected (4 nl/sec, 600 nL/hemisphere). For electrophysiology experiments, holes were drilled with a 0.9 mm burr above the ACC (A/P:±1.7 mm, M/L:+0.3 mm), LC (A/P -5.4mm, M/L: +0.9mm), and lambda.

Two stereotrodes (Cat #8245-M, Pinnacle Technologies) attached to a head-mount, with two wires soldered to bone screws were implanted unilaterally in ACC (A/P:+1.7mm, M/L:+0.3 mm, D/V: -1.7) and LC (A/P:-5.4, M/L:+0.9, D/V:− 3.5).

### Behavioral training and scoring

The rCPT training protocol was based on a previously described protocol [39]. Briefly, mice were trained in Bussey-Saksida mouse touchscreen chambers (Lafayette Instruments). Training stage details are provided in supplement. Performance scoring parameters were similar to those described by <u>[20]</u> and <u>[39]</u>. Briefly, to assess attention performance during Stage 3, we calculated the discrimination index **d’**, which is a measure of sensitivity bias (perceptual discriminability between S+ and S−) and **c**, which is a measure of response bias (willingness to make responses).

### Immunohistochemistry for c-Fos

Coronal sections were incubated in 1:1000 anti-c-Fos antibody (SySy; cat # 226003), 1:1000 goat anti-rabbit AlexaFluor 555 (Sigma), and 1:5000 DAPI (Sigma). Images were visualized at 40x.

### In vivo electrophysiology and analysis

LFPs were recorded: 1) After mice reached performance criteria (≥ 60 hits/session) during Stage 2; 2) during the first Stage 3 session (Stage 3-early); and 3) after mice reached performance criteria during Stage 3 (Stage 3-late). LFPs were sampled at 2 kHz, and multitaper spectral analysis was used for power density estimation and phase coherence (http://chronux.org[37]). For phase-amplitude coupling, values for frequency pairs were extracted via Morlet wavelet convolution. Phase and amplitude values were binned and mean amplitude values were normalized by dividing each bin value by the summed value over all bins. A modulation index value was derived by calculating the Kullback-Leibler distance between the observed phase-amplitude distribution and a uniform (null) phase-amplitude distribution <u>[40]</u>. For Granger causality analysis, we performed multiple lagged autoregressions on LFP traces with the model order for each set of autoregressions determined by the Bayes’ information criteria. Values for directionality were normalized into a lead index value.

## RESULTS

### LC-ACC circuit is recruited during rCPT performance

The ACC is necessary for rCPT performance, but how ACC circuits control distinct aspects of behavior, and which circuits provide neuromodulatory control over the ACC to regulate this behavior are not known. To better understand ACC-associated circuits involved in rCPT behavior, we asked which afferent projections to the ACC are selectively activated during task-related behavior. To answer this question, we used a behavioral circuit-tracing approach that allowed us to map recruitment of neurons projecting to the ACC across two stages of rCPT training (Stage 2 vs. Stage 3) with different cognitive demand requirements (Fig. 1A, B). We chose several candidate regions that have been implicated in attention-related behavior and that have direct projections to the ACC, including the ventral hippocampus (vHC), medio-dorsal thalamus (MD), and the locus coeruleus (LC). We injected AAVrg-CAG-tdTomato into the ACC to label these afferent projections. Following viral labeling, mice were trained until they achieved asymptotic performance on Stage 2 of the rCPT (Supplementary Fig. 1). Half of the mice were further trained on Stage 3 (Stage 3 group), while the other half were continuously run on Stage 2 (Stage 2 group). Once a mouse in the Stage 3 group reached performance criteria, that mouse and a yoked mouse from the Stage 2 group were killed two hours following a behavioral session. Brains were then collected for c-Fos immunohistochemistry and imaging. Double labeling for c-Fos and tdTomato was quantified to determine the proportion of neurons projecting to the ACC from the chosen regions that were activated during different stages of rCPT training (Fig. 1C and D). Within these brain regions, the LC contained the highest percentage of c-Fos+/tdTomato+ neurons across both groups (Stage 2: MD= 11.3% ± 11.8%; vHC= 3.17% ± 5.5%; LC= 28.2% ± 31.4%; Stage 3: MD= 14.9% ± 13.8%; vHC= 2.38% ± 2.07%; LC= 66.4% ± 29.2%; means ± standard deviation) (Fig. 1C and Supplementary Fig. 2). Moreover, more ACC-projecting LC neurons were recruited during Stage 3, when cognitive demand is higher, compared to yoked Stage 2 controls (*t*(22) = -3.0855, *p* = 0.005; Fig. 1C-D) This effect was not observed in either the MD (*t*(7) = -0.42438, *p* = 0.684), or the vHC (*t*(4) = 0.23309, *p* = 0.8271) (Supplementary Fig. 2).

**Figure 1.**
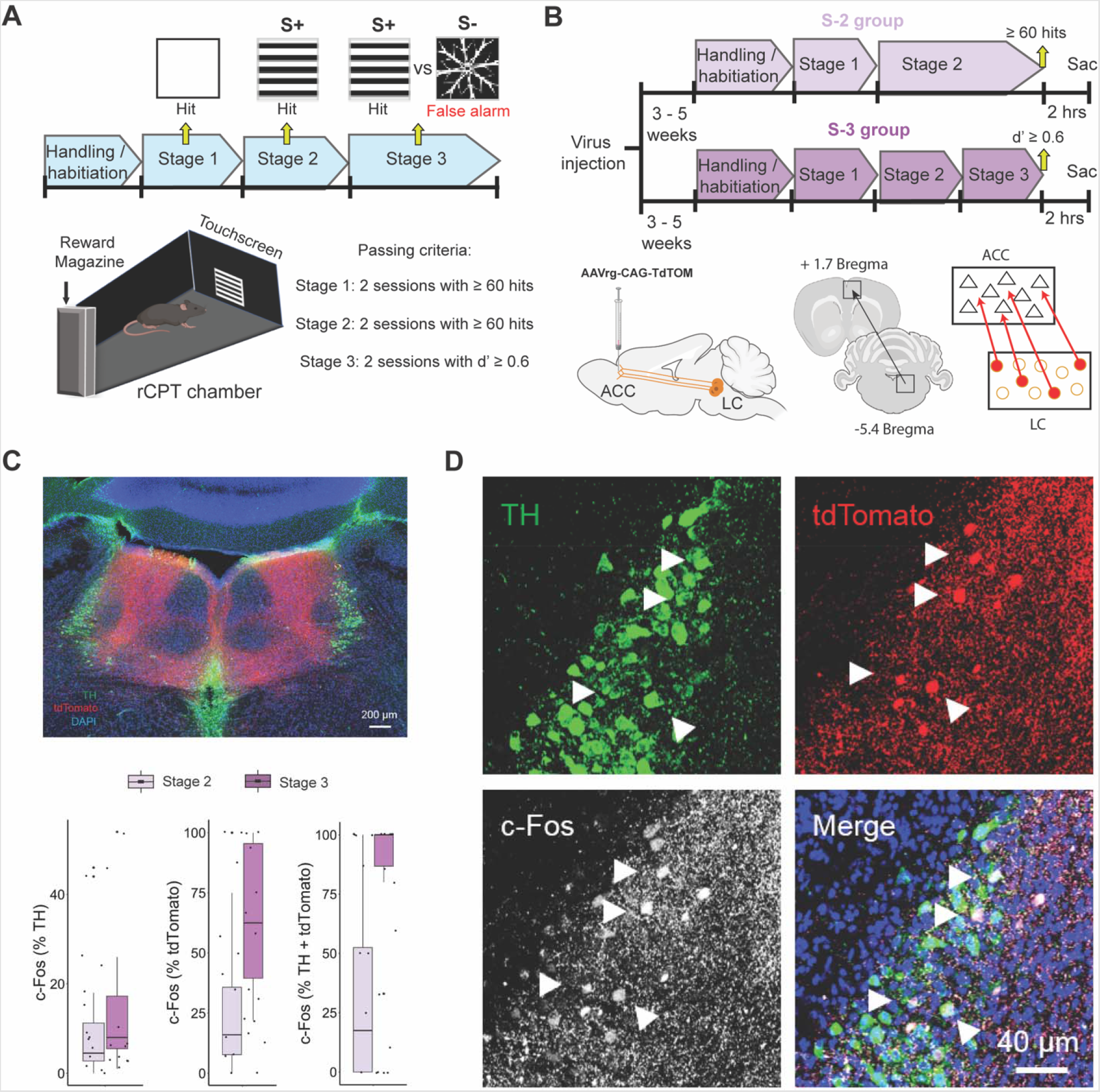
rCPT training recruits ACC-projecting LC neurons. A) Timeline of rCPT training (top). Arrows indicate the type of stimuli presented across training stages and the response when interacting with the stimulus. Mice are moved to the next training stage when they reach performance criteria (bottom right) in two consecutive sessions. Schematic of the touchscreen chambers (bottom left) depicting a stimulus presentation during the rCPT at the center of the touchscreen located in front of the chamber and the reward magazine in the back. B) Timeline of c-Fos experiments (top) and schematics of viral injections designed to label ACC-projecting LC neurons (bottom). C) Low magnification confocal image of an LC coronal slice immunostained against TH (green), TdTomato (red), c-Fos (white), and DAPI (Blue) (top), and boxplots summarizing the quantification c-Fos in TH+ (left), tdTomato+ (middle) and TH+/tdTomato+ (right) neurons in the LC. D) High magnification confocal images of TH, tdTomato and c-Fos expression in LC neurons. Arrows show examples of ACC-projecting (tdTomato+) and NE (TH+) neurons in the LC that co-express c-Fos following the last session of rCPT training.

### Cognitive demand and proficiency during rCPT training increase power of low frequency bands within ACC and LC LFPs

To better understand the role of LC-ACC circuitry in sustained attention during rCPT performance, we recorded simultaneous local field potentials (LFPs) within the ACC and LC at three timepoints: Stage 2, Stage 3-early, and Stage 3-late (Fig. 2). At the Stage 2 time point, mice have demonstrated proficiency in detecting stimuli presented on the touchscreen and responding appropriately with a nose poke (Supplementary Fig. 4A). In Stage 3-early, which is the first Stage 3 session following advancement from Stage 2, mice are required to not only detect stimuli on the touchscreen, but to discriminate between the S+ and S- and respond accordingly. Performance, measured as d’, is lower in Stage 3-early compared to Stage 3-late, which is the last Stage 3 session, during which overall performance reached the established criteria. As expected, mice in the Stage 3-late group had better task performance than mice in the Stage 3-early group, as reflected by a significantly higher mean d□ score (*t*(19) = -9.4951, *p* < 0.001) (Supplementary Fig. 4A and B). Despite better performance, mice in the Stage 3-late group did not tend to change their overall response strategy (as reflected in their mean c score), preferring to maintain a fairly conservative response bias across both Stage 3-early and Stage 3-late sessions (*t*(19) = 0.20951, *p* = 0.8363) (Supplementary Fig 4C). Latency to the reward trough also did not differ between Stage 3-early and Stage 3-late sessions. However, there was a significant difference in the response latency between hits and false alarms, with mice responding significantly faster during false alarms than hits (*F*(1,38) = 8.364, *p* = 0.0063, main effect of response type for session x response type mixed-factorial ANOVA) (Supplementary Fig. 4D). These data show that mice improve their ability to discriminate between the S+ and the S-over time without changing their response bias while false alarms are associated with an impulsive motor pattern.

**Figure 2.**
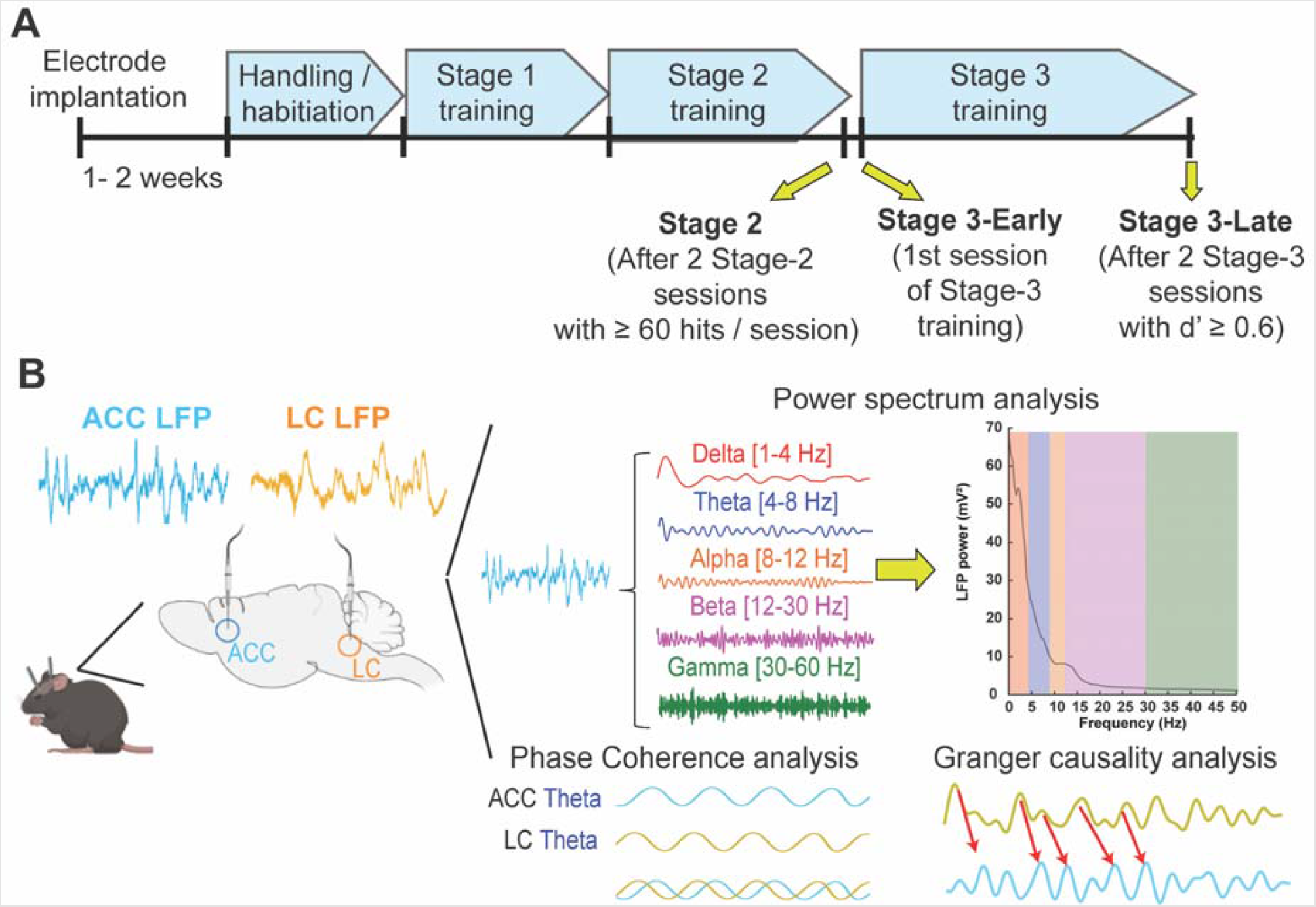
LFP recording sessions within rCPT training stages. A) Timeline of rCPT training showing the three recording sessions: Stage 2, Stage 3-Early, and Stage 3-Late. B) Schematic of depth electrode placements within the ACC and the LC (left) and summary of LFP data analysis (right)

To understand the contributions of LC-ACC circuitry to rCPT learning and performance, we analyzed ACC and LC LFPs in the 4-second time window surrounding mouse responses (screen touches) to the S+ (hits) and S-(false alarms) to identify electrophysiological correlates of performance across recording sessions. We further split this time window into two distinct timepoints centered around the screen nosepoke: pre-screen touch (from -2 to 0 seconds), and post-screen touch (from 0 to 2 seconds). We first analyzed amplitude (power) within the ACC LFP during these timepoints, and found differences in lower-frequency bands during hits and false alarms (Fig. 3A). Specifically, we found no significant difference in either delta (1-4 Hz), or theta (4-12 Hz), power (*p* > 0.05 for session x response type mixed-factorial ANOVA) during pre-screen touch, but found that both delta (*F*(1,38) = 4.54, *p* = 0.0396) and theta (*F*(1,38) = 6.115, *p* = 0.018) power were significantly higher during post-screen touch hits vs. false alarms in both Stage 3-early and Stage 3-late sessions (main effect of response type for session x response type mixed-factorial ANOVA) (Fig. 3 B-F). When analyzing the same frequency bands from the LC LFP, a different pattern emerged; namely, post-screen touch, but not pre-screen touch, theta and delta power differed between hits and false alarms for Stage 3-late sessions, but not Stage 3-early sessions (*F*(1,38) = 5.107, *p* = 0.0297, session x response type interaction) (Fig. 4). These data suggest a disparity between the ACC and LC with regard to task engagement during rCPT learning, with the ACC differentiating between hits and false alarms (the S+ and S-) regardless of task proficiency, and the LC differentiating between the two response types only when the task is well-learned.

**Figure 3.**
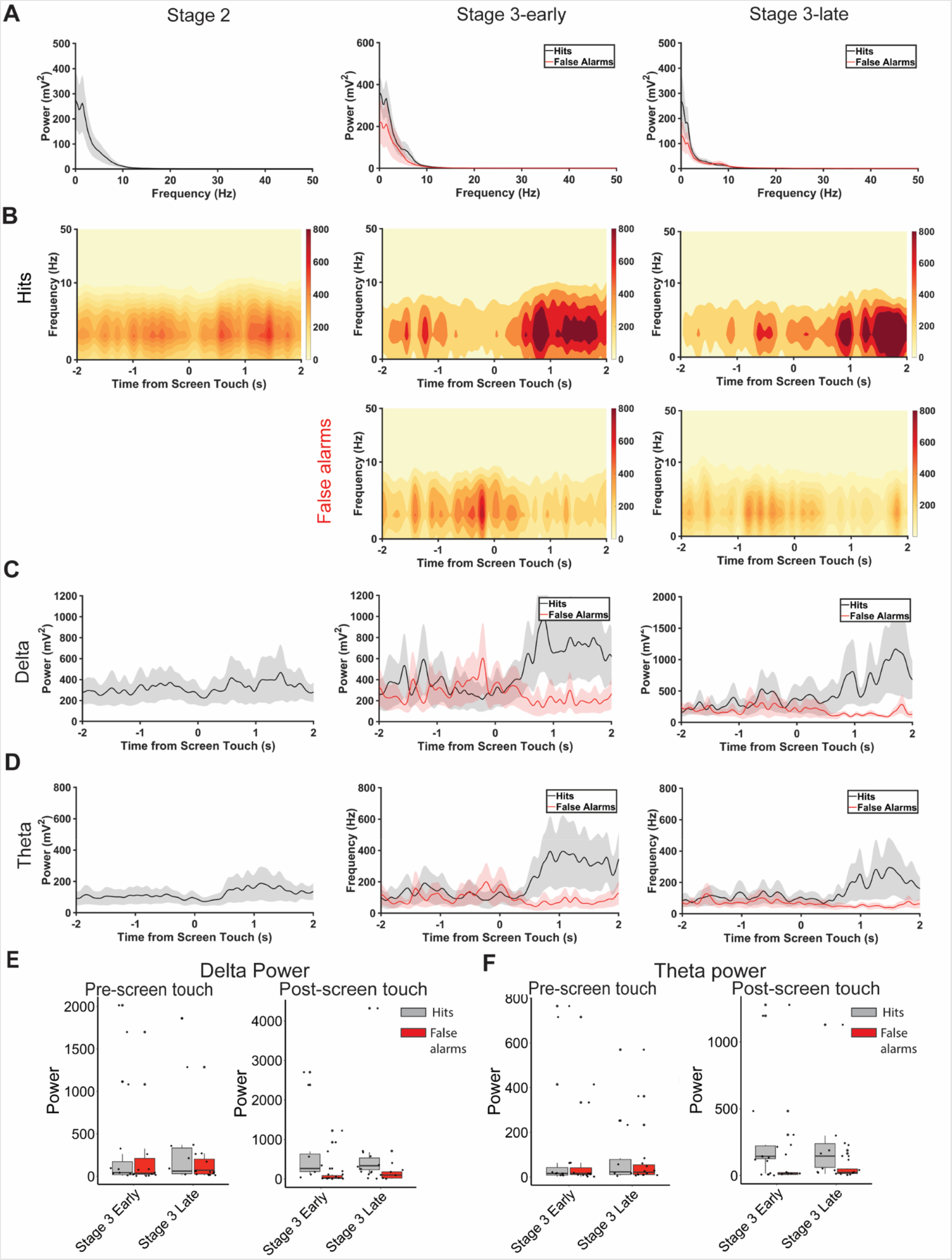
Correct responses (hits) during Stage 3 of rCPT training increase delta and theta power within the ACC. A) Power spectral density of ACC electrode during hits (black) and false alarms (red) in Stage 2 (left), Stage 3-early (middle), and Stage 3-late (right) recording sessions. B) Spectrograms of ACC electrode surrounding hits (top) and false alarms (bottom) in Stage 2 (left), Stage 3-early (middle), and Stage 3-late (right) recording sessions. C-D) ACC delta (C) and theta (D) power across time surrounding hits and false alarms in Stage 2 (left), Stage 3-early (middle), and Stage 3-late (right) recording sessions. E) Boxplots summarizing changes in ACC delta power before (left) and after (right) screen touch during hits and false alarms. F) Boxplots summarizing changes in ACC theta power before (left) and after (right) screen touch during hits and false alarms.

**Figure 4.**
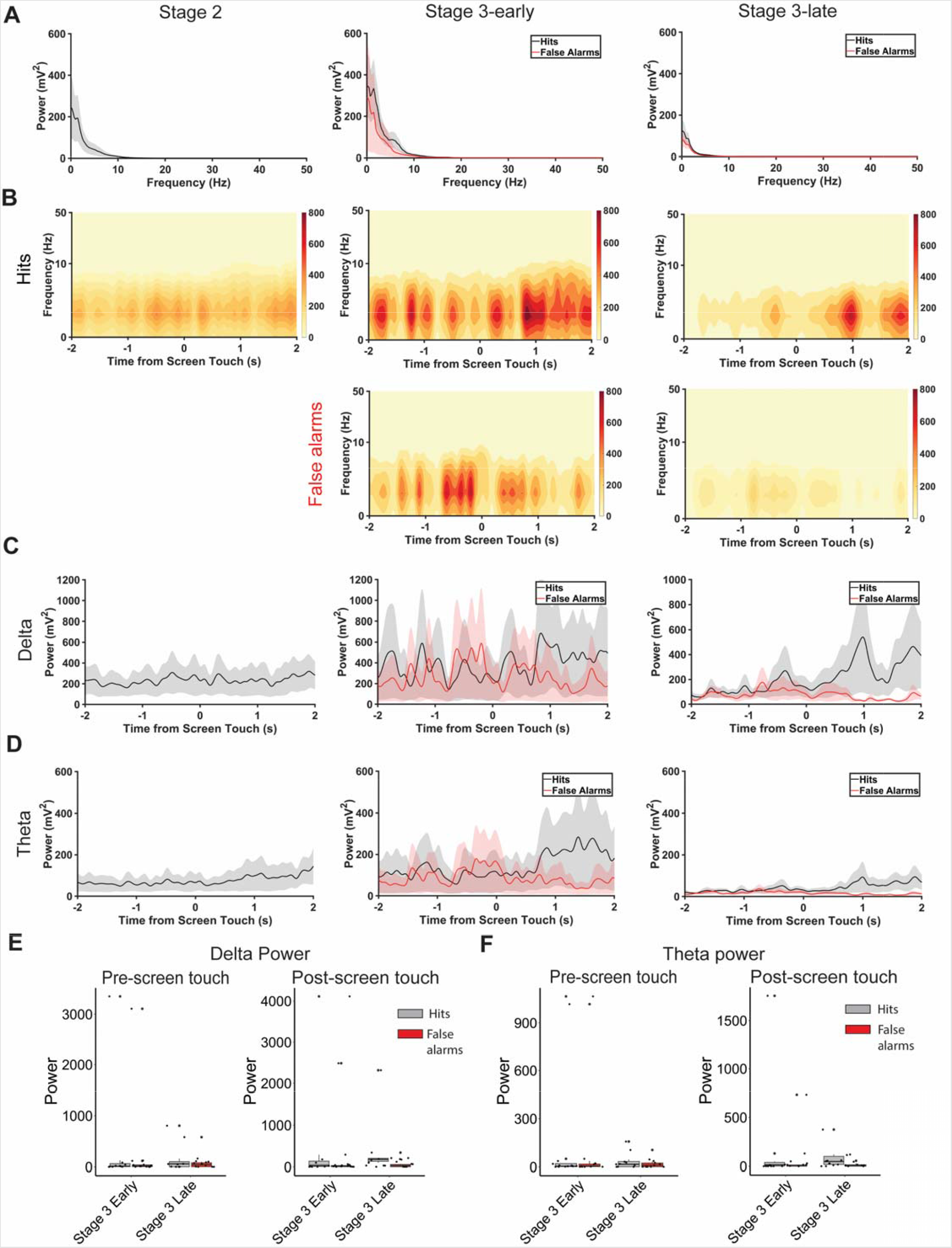
Correct responses during Stage 3-late of rCPT training increase delta power within the LC. A) Power spectral density of LC electrode during hits (black) and false alarms (red) in Stage 2 (left), Stage 3-early (middle), and Stage 3-late (right) recording sessions. B) Spectrograms of LC electrode surrounding hits (top) and false alarms (bottom) in Stage 2 (left), Stage 3-early (middle), and Stage 3-late (right) recording se sions. C-D) LC delta (C) and theta (D) power across time surrounding hits and false alarms in Stage 2 (left), Stage 3-early (middle), and Stage 3-late (right) recording sessions. E) Boxplots summarizing changes in LC delta power before (left) and after (right) screen touch during hits and false alarms. F) Boxplots summarizing changes in LC theta power before (left) and after (right) screen touch during hits and false alarms.

To further assess relationships between activity in the ACC and behavior during CPT learning, we analyzed the degree to which the phase of delta and theta oscillations modulates the amplitude of gamma oscillations during screen touch. Phase-amplitude coupling between oscillations in different frequency bands has been linked to numerous cognitive functions, including attention [41–43]. We found a selective effect of response type (hits vs. false alarms) on delta-slow gamma (30-55 Hz) phase-amplitude coupling in the ACC LFP (*F*(1,38) = 4.451, *p* = 0.0415, main effect of response type for session x response type mixed-factorial ANOVA) that was not present for coupling between other frequency bands (delta-fast gamma (65-120 Hz), theta-slow gamma, or theta-fast gamma) (Supplementary Fig. 5). Delta-slow gamma coupling and delta power therefore increase in tandem during hits, regardless of which session (Stage 3-early or Stage 3-late) the mouse is performing.

### The LC and ACC dynamically interact during rCPT training and proficient task performance

To investigate how the LC and ACC interact during rCPT performance, we assessed the degree to which LFPs in the two brain regions phase-locked with one another across different stages. Phase coherence in the delta frequency band was significantly higher during hits vs. false alarms during both Stage 3-early and Stage 3-late sessions (*F*(1,38) = 7.77, *p* = 0.0083, main effect of response type for session x response type mixed-factorial ANOVA) (Fig. 5A and B). Interestingly, although theta phase coherence did not significantly differ between response types, it did significantly decrease from Stage 3-early to Stage 3-late sessions (*F*(1,38) = 8.836, *p* = 0.0051, main effect of session for session x response type mixed-factorial ANOVA) (Fig. 5A and B), indicating that distinct pieces of information (the mouse’s response vs. the session the mouse is performing) may be routed between the LC and ACC in disparate frequency bands.

**Figure 5.**
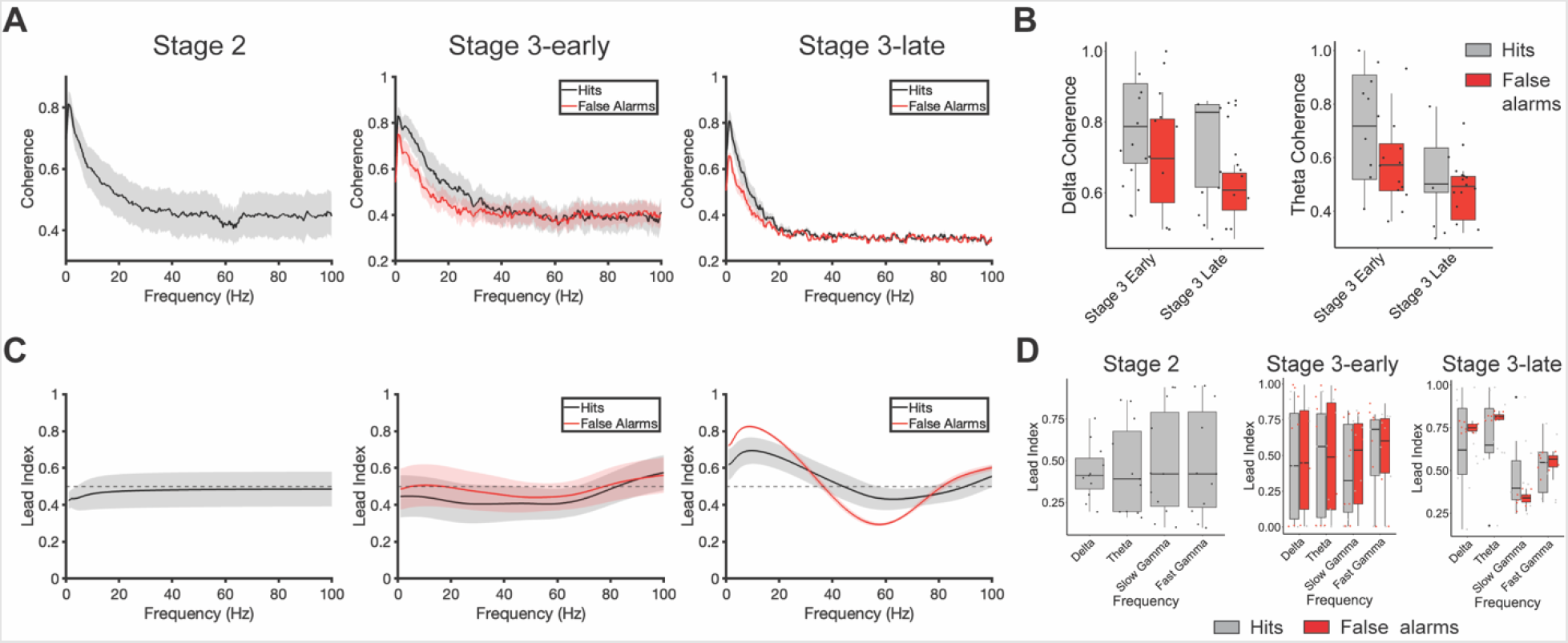
LC leads ACC in theta and beta frequencies during hits in Stage 3-late. A) Coherence analysis of LC and ACC LFPs across Stage 2 (left), Stage 3-early (middle), and Stage 3-late (right) recording sessions. Stage 3-late false alarms exhibit lower coherence within low frequency bands (0-8 Hz) than Stage-3 hit. B) Boxplots summarizing coherence across LC and ACC LFPs within delta (0-4 Hz; left) and theta (4-8 Hz; right) bands during hits (gray) and false alarms (red). C) Granger causality index across recording sessions. Lead index is normalized where values higher than 0.5 (dotted line) indicate LC leading ACC, while values below 0.5 indicate ACC leading LC. Only Stage 3-late shows directionality where LC leads ACC in theta/beta frequencies while ACC leads LC in gamma frequencies during false alarms. D) Boxplot summarizing lead index across recording sessions.

Since the LC-ACC circuit is bi-directional with neurons in the LC sending axons to the ACC, and the LC additionally receiving direct input from ACC neurons, we investigated whether the LC conveys information to the ACC, or vice versa, during distinct rCPT training stages. To infer directionality, we used Granger causality to determine how the LFP in one brain area predicted future changes in the LFP in the other brain area. We then normalized Granger causality values into a “lead index” variable, which allowed us to determine which brain region led the other within defined frequency bands (lead index values > 0.5 indicate that the LC LFP predicts the ACC LFP, and lead index values < 0.5 indicate that the ACC LFP predicts the LC LFP). During Stage 2, there was no clear pattern of directionality across delta, theta, slow gamma (30-55 Hz), or fast gamma (65-120 Hz) frequency bands (all lead index values not significantly different from 0.5, *F*(3,40) = 0.088, *p* = 0.966, one-way ANOVA across frequency bands with lead index as the dependent variable) (Fig. 5C and D). A similar pattern emerged during Stage 3-early sessions for both hits and false alarms (*p* > 0.05 for frequency band x response type mixed-factorial ANOVA) (Fig. 5C and D). During Stage 3-late sessions, however, the LC LFP led the ACC LFP in lower frequency bands (delta and theta) during hits (Fig. 5C and D). During false alarms, the LC LFP also led the ACC LFP in these frequency bands, but the ACC LFP additionally led the LC LFP in the slow gamma frequency band (*F*(3,72) = 19.712, *p* < 0.001, main effect of frequency band, and *F*(3,72) = 4.645, *p* = 0.033, frequency band x response type interaction for frequency band x response type mixed-factorial ANOVA) (Fig 5C and D), suggesting that communication from the LC to the ACC is a hallmark of proficient task performance generally, but that the ACC additionally communicates back to the LC during errors.

## DISCUSSION

The LC plays a major role in general arousal and vigilance [44–46]; for review [47]. Phasic activation of LC neurons is associated with both sleep-to-wake transitions [48,49], and orienting the response to salient or unexpected sensory stimuli [50] for review [51]. NE-producing neurons in the rodent LC send efferents to many cortical areas, including the ACC [52], and arousal-related patterns of neuronal activity in the ACC are linked with phasic LC activity [53,54]. The “adaptive gain theory” of LC function posits that enhanced attention is, in part, a direct result of the arousal-inducing properties of phasic activity in the LC [55,56]. Supporting this idea, lesions of LC-NE efferents to the frontal cortex impair attentional performance in rodents [57,58] and non-human primates [59]; for review [60], and catecholaminergic signaling in the ACC regulates behavior in attention tasks [61,62], including the rCPT [24,63,64]. Our results further support the notion that communication between the LC and the ACC regulates successful stimulus discrimination and discrimination-guided behavior in mice. Specifically, viral labeling and immunohistochemical analysis showed that LC neurons that target the ACC are selectively activated after mice learn to asymptotically perform the touchscreen-guided rCPT. We investigated recruitment of attention-related immediate early gene expression in other ACC input regions, including the MD and vHC. In contrast to the LC, we found no evidence that ACC inputs from the MD and vHC are activated during rCPT performance. The MD is the major thalamic input to the frontal cortex [65,66], and is also heavily involved in attentional processes [67–69]. That LC-ACC projection neurons, but not MD-ACC projection neurons, were recruited during the rCPT suggests that these two pathways functionally contribute to different aspects of attention in rodents. These findings are in line with the theory that LC-mediated NE release in the ACC promotes sustained attention.

To investigate the roles of the ACC and LC in sustained attention, we simultaneously recorded LFPs in these brain areas during rCPT performance. We found that activity in both regions distinguishes between correct and incorrect responses (e.g. hits and false alarms). These patterns of activity, however, are only present in the LC when the rCPT is well-learned, suggesting that the LC encodes the attention-related task rule. Specifically, amplitude in the delta and theta frequency bands is increased during hits (directly after the mouse pokes the screen), supporting a role for the LC in the exploitation of correct responses. This result is perhaps surprising given that both increased activity in the LC [70], and increases in pupil diameter [71,72] are correlated with uncertainty during decision-making [73]. Both optogenetic [74], and chemogenetic [75], stimulation of LC neurons, however, accelerates learning a new rule (exploitation), rather than accelerated disengagement from an old role (exploration), indicating that the LC promotes cognitive flexibility by facilitating adaptation to new behavioral policies. The LC may promote both exploration and exploitation via tonic and phasic firing of LC neurons, respectively [55,76,77]; if this is the case, then activity in the LC that leads to phasic spiking may also underlie increases in low-frequency oscillations in the LC LFP during well-learned attention-guided behavior. A limitation of this study is that LFPs were only recorded during the first rCPT Stage 3 session (Stage 3-early), and the final rCPT Stage 3 session (Stage 3-late), which precludes analyzing how brain-behavior relationships shift during the entire learning process. Understanding how delta and theta activity in the LC LFP develop as a function of learning will be informative for understanding the relationship between the LC and exploration-exploitation during this task.

In contrast to activity in the LC, delta and theta power in the ACC LFP distinguishes between hits and false alarms even when mice have not yet learned the task rule. Indeed, response time differences in this study demonstrate a dissociation between visual discrimination and behavioral choice, indicating that mice discriminate between the target and non-target well in advance of learning to respond differently to the two [78]. Increased ACC power tracks this visual discrimination across orthogonal learning states, consistent with a role for the ACC in attention-related sensory processing [79,80] and decision making during both periods of certainty and uncertainty [81,82]. Concurrently with increased delta power, phase-amplitude coupling between delta and slow-gamma oscillations is also increased during hits, regardless of session. Phase-amplitude coupling in the ACC is linked to visual cue detection [42], reaction time [43], and stimulus orientation [83] during attentional tasks, signaling a role for the ACC in attention-guided sensory discrimination. The dissociation between LFP activity in the LC and ACC in this study suggests that these two brain areas process distinct aspects of experience during correct and incorrect choices, and argues against the possibility that motor behavior or reward consumption (the temporal onset of power differences between hits and false alarms is highly aligned to screen touch, occurring before the mouse reaches the reward port) is driving this dissociation, as would be expected if both brain areas responded similarly during Stage 3-early and Stage 3-late sessions.

LC-NE neurons make synaptic contact with [84–87], and also receive synaptic contact from the ACC [88,89], suggesting that these two brain areas may functionally communicate during the rCPT. To test this, we analyzed phase coherence (the degree to which phases within a defined frequency band are aligned on successive cycles) between LFPs in the LC and ACC during Stage 3-early and Stage 3-late sessions. Coherence in the delta frequency band was highest during hits, regardless of session type, and coherence in the theta frequency band was highest during Stage 3-late sessions, regardless of response, signaling a dissociation between the type of information that is conveyed by these two oscillations. To uncover directionally-specific (i.e., from LC-to-ACC, or vice versa) patterns of communication between the two brain areas, we used Granger causality, and found that LC-to-ACC directionality in the delta and theta frequency bands was highest during Stage 3-late sessions. This directionality did not reach statistical significance during either Stage 2 or Stage 3-early sessions, indicating that phase coherence during Stage 3-early is a product of bi-directional transmission between the LC and ACC. LC-to-ACC low-frequency directionality during Stage 3-late sessions was observed during both hits and false alarms, implicating this pattern of directionality in attention-based discrimination performance (i.e., in overtrained animals), rather than attention-based discrimination learning. Directionality from the ACC-to-LC, however, did discriminate between hits and false alarms - specifically, ACC-to-LC directionality in the slow gamma frequency band was higher during false alarms, suggesting that the ACC conveys an error signal to the LC during higher periods of certainty in attention tasks. This finding represents a potential behavioral correlate to data in anesthetized rats showing that gamma power in the ACC precedes low-frequency activity in the LC [90], and provides experimental data to support a recently proposed computational model where ACC inputs to the LC encode prediction errors [91].

In summary, our findings reinforce the well-known roles of the LC and ACC in attention, and further demonstrate multiplexed relationships between specific patterns of oscillatory synchrony between the two brain regions and different stages of learning in a translationally-relevant attentional discrimination task. That communication between the LC and ACC is routed through distinct frequency bands at different timepoints is perhaps unsurprising in light of the complex relationships between multi-unit activity in these two areas, even when animals are anesthetized [92,93]. The results described here could be compared with neural responses during human CPTs to identify a translational benchmark for circuit function in patients with attentional deficits. This may be useful because attention deficits related to LC-ACC function are supported across multiple complex brain disorders, including major depressive disorder [94], post-traumatic stress disorder [95], schizophrenia [96]. Hence, establishing biomarkers of function in this circuit during attentional tasks is of significant value.

## FUNDING

This work was supported by internal funding from the Lieber Institute for Brain Development, and the National Institute of Mental Health (R01MH105592 to KM and R56MH126233 to GVC and KM).

## ACKNOWLEDGEMENTS

We thank members of the Martinowich and Carr laboratories for helpful comments and suggestions. We also thank Aimee Ormand and Deveran Manley for assistance with animal care.

## AUTHOR CONTRIBUTIONS

Conceptualization: HLH, GVC, KM

Methodology: HLH, JMM, JMB, SA, ACD, YL

Validation: JMB, LO Formal Analysis: HLH, SA

Investigation: HLH, SA, JMB, JMM, LO

Data Curation: HLH, SA, JMB

Writing-original draft: HLH, SA, JMB, GVC, KM

Writing-review and editing: HLH, SA, JMB, GVC, KM

Visualization: HLH, SA, JMB

Supervision: HLH, JMB, KM

Project administration: JMB, KM, GVC

Funding Acquisition: GVC, KM

## COMPETING INTERESTS

GVC is a scientific advisor for LongTermGevity, Inc. and owns stock options in the company. LongTermGevity, Inc. was not involved in the funding, design, or execution of these studies. No other authors have financial relationships with commercial interests, and the authors declare no competing interests.

