## Supplemental Information for "Electrophysiological correlates of attention in the locus coeruleus - anterior cingulate cortex circuit during the rodent continuous performance test"

**Supplemental Methods**

***Animals***

Male C57BL/6J mice (Strain#: 000664) were purchased directly from The Jackson Laboratory (Bar Harbor, ME, USA), and were eight to ten weeks old at the start of all experiments. Upon arrival in the facility, mice were acclimated to the colony room for 72 h prior to starting experimental procedures. Mice were group-housed (4/cage) until electrode implantation (see surgical procedure) and then double-housed for the remainder of the experiment. Mice were housed in disposable polycarbonate caging (Innovive, San Diego, CA, USA) and maintained on a reverse 12/12 light/dark cycle (lights on at 19:00 hours/lights off at 07:00 hours). Mice had ad libitum access to Teklad Irradiated Global 16% Protein Rodent Diet (#2916; Envigo, Indianapolis, IN, USA) in the home cage until the food restriction protocol was initiated. Water was available in the home cage ad libitum throughout all experiments. Behavioral testing was conducted Monday-Friday during the dark phase (10:00-17:00 hours). All experiments and procedures were approved by the Johns Hopkins Animal Care and Use Committee and conducted in accordance with the Guide for the Care and Use of Laboratory Animals.

***Food restriction protocol***

Mice were handled and weighed for at least two consecutive days before starting the food restriction protocol. This protocol entailed restricting available food to 2.5 g of chow per mouse per day, and weighing mice daily to ensure that weight was maintained at 85-90% of their predicted free-feeding weight based on average growth-curve data for the strain (The Jackson Laboratory, Bar Harbor, ME). To familiarize the mice to Nesquik® strawberry milk (Nestlé, Vevey, Switzerland), which was used as reward during rCPT training, a 4x4 inch weighing plate (VWR, Radnor, PA, USA) containing ~2 mL of strawberry milk was introduced to the home cage for three consecutive days. The weighing plate was left in the cage until all mice had sampled the strawberry milk.

***Surgical procedures***

Mice were anesthetized with isoflurane (induction: 2-4% in oxygen, maintenance: 1-2%) and then secured to a stereotaxic frame. The top of the skull was exposed by an incision along the midline of the scalp, Bregma and Lambda were identified, and the head was leveled to ensure the skull was flat. For immunohistochemistry experiments, small holes were drilled with a 0.9 mm burr (Fine Science Tools., Foster city CA) above the ACC (A/P: ±1.7 mm, M/L: + 0.3 mm), and a retrograde virus expressing tdTomato (AAVrg-CAG-tdTomato, Addgene catalog # 59462-AAVrg) was injected at a rate of 4 nl/sec, and a total volume of 600 nL per hemisphere. For electrophysiology experiments, small holes were drilled with a 0.9 mm burr (Fine Science Tools., Foster city CA) above the ACC (A/P: ±1.7 mm, M/L: + 0.3 mm), LC (A/P: -5.4 mm, M/L: + 0.9 mm), and lambda. Two stereotrodes (Cat #8245-M, Pinnacle Technologies) attached to a head-mount, along with two wires that were soldered to two bone screws (Fine Science Tools) were slowly implanted unilaterally into the ACC (A/P: +1.7 mm, M/L: + 0.3 mm, D/V: -1.7) and LC (A/P: -5.4, M/L: +0.9, D/V: −3.5). Head-mount ground screws were implanted contralaterally to the ACC electrode and directly above lambda. The entire head mount was secured to the skull using dental acrylic (Ortho-jet, Lang Dental manufacturing Co.). Following surgery, mice received a subcutaneous injection of a local anesthetic (Bupivacaine) and recovered on a warm heating pad before being transferred back to the colony room. For the following three days after surgery, mice received Meloxicam injections (20 mg/kg) to relieve pain, and were monitored for any abnormal behavior. Mice remained in the colony room for at least 1 week to allow full recovery before initiating behavioral training.

***Behavioral training***

*Habituation*: Mice received two consecutive habituation sessions (20 min each) to acclimate them to the touch screen chambers (Lafayette Instruments, Lafayette, IN, USA). During habituation sessions, 1 mL of strawberry milk was placed into the reward tray. The screen was responsive to touch, but touches were not rewarded. Mice then were moved to rCPT training if in at least one session the mouse had consumed the strawberry milk.

*Rodent Continuous Performance Test (rCPT):* The rCPT training protocol was based on a previously described protocol[[1]](https://sciwheel.com/work/citation?ids=12028668&pre=&suf=&sa=0). Briefly, mice were trained in Bussey-Saksida mouse touchscreen chambers (Lafayette Instruments, Lafayette, IN, USA) connected to a computer using ABET II software (Campden Instruments, Loughborough, UK).

*Stage 1:* Mice were trained Monday-Friday during 45 min sessions to respond to a visual stimulus (white square) presented at the center of the touch screen. The stimulus was displayed for 10 sec (stimulus duration, SD), where a touch in the center of the screen produced delivery of ~20 μL of strawberry milk in the reward tray located on the opposite side of the touch screen chamber. Following SD, a 0.5 sec limited hold (LH) period was given during which the screen was blank, but a touch would still yield a reward. Upon interacting with the stimulus, a one sec tone (3 kHz) was delivered, the reward tray was illuminated to signal reward delivery, and the schedule was paused until a head entry into the reward tray was detected by an IR beam. Then, a 2 sec intertrial interval (ITI) would begin before the subsequent trial started. If the mouse did not interact with the stimulus during the SD or LH, an ITI would start, and the next trial would follow. The criterion for a mouse to advance to the next stage was to obtain at least 60 rewards per session in two consecutive sessions.

*Stage 2:* In Stage 2, a target stimulus (S+) was introduced. The S+ consisted of a square with either horizontal or vertical black and white bars that replaced the white square at the center of the screen. Sessions were still 45 min long and each mouse was assigned either horizontal or vertical oriented S+ for the remaining sessions of the experiment. The S+ assignment was counterbalanced for each group. During Stage 2, the SD was 5 sec, and LH was 5.5 sec. Interaction with S+ (Hit) during SD + LH resulted in reward delivery. Once a mouse obtained at least 60 hits/session in two consecutive sessions, an *in vivo* electrophysiological recording session (see *in vivo* electrophysiology) followed the day after. A mouse was moved to the next stage if it obtained at least 60 hits in the recording session. If a mouse failed to obtain at least 60 hits in the recording session, the recording session was repeated until the mouse obtained at least 60 hits.

*Stage 3:* In Stage 3, a non-target stimulus (S-) consisting of a snowflake shaped stimulus presented at the center of the screen was introduced. On each trial, the probability of S+/S- was 50%/50%. The SD was 3 sec, the LH was 3.5 sec, and the ITI length was either 2 or 3 sec in length (randomized ITI duration during trials). Similar to Stage 2, screen touches during S+ (hit) yielded a reward, but screen touches during S- did not (false alarm). A false alarm resulted in the beginning of the ITI followed by a correction trial. In correction trials a S- was presented again. If another false-alarm occurs, a new correction trial starts until the mouse doesn’t interact with the S- (correct rejection). To determine attention performance during Stage 3, a discrimination index (d’, see behavioral scoring) was used. Mice were trained in Stage 3 for at least seven sessions, and until they had a d’ score of 0.6 or higher in the last two consecutive sessions. The experiment ended with a recording session where mice were required to obtain a d’ of 0.6 or higher with at least 10 false alarms. In the scenario where mice didn’t reach the criteria during the recording session, the session was repeated until mice reached the criteria.

Behavioral scoring:

Behavioral databases containing the timestamps from stimuli presentation, hits, false alarms, latency to response, etc. were retrieved from ABET II (Lafayette Instruments, Lafayette, IN, 204 USA) and Whisker server (Cambridge University Technical Services, UK). Then behavioral data was analyzed in Excel to ascertain performance parameters. Performance scoring parameters were similar to those described by[[2]](https://sciwheel.com/work/citation?ids=2724342&pre=&suf=&sa=0) and[[1]](https://sciwheel.com/work/citation?ids=12028668&pre=&suf=&sa=0). Briefly, to assess attention performance during Stage 3 training, we calculated discrimination index d’,which is a measure of sensitivity bias (refers to the perceptual discriminability between the S+ and S−) and c which is a measure of response bias (refers to the criterion or willingness to make responses) with the following formulas:

d’ = z(Hit rate) - z (False alarm rate)

c = -(z(Hit rate) + z (False alarm rate))

2

Whereas:

Hit rate (HR) = Hits / Hits + misses

False Alarm rate (FAR) = False alarms / False alarms + correct rejections.

***Immunohistochemistry for c-Fos***

Coronal sections (50 μm) were cut on a sliding microtome (Leica) with attached freezing stage (Physitemp), washed in 5% Tween-80 in 1× PBS, and incubated in blocking solution (0.5% Tween-80, 5% normal goat serum in 1× PBS) with agitation for 6–8 h. The sections were then incubated in 1:1000 anti-c-Fos antibody (SySy; cat # 226003) in blocking solution overnight at 4 °C with agitation. The following day, the sections were washed, incubated in 1:1000 goat anti-rabbit AlexaFluor 555 (Sigma) in blocking solution for 2 h with agitation, washed again, and incubated in 1:5000 DAPI (Sigma) in 1x PBS for 20 min. Fos protein fluorescence was visualized on a Zeiss LSM 700 confocal microscope with a 40x oil-immersion lens.

***In vivo electrophysiology and analysis***

Local field potentials (LFPs) from the ACC and the LC were recorded using Sirenia acquisition software (Pinnacle Technology Inc, USA) at three different timepoints of behavioral training: 1) After mice reached performance criteria (≥ 60 hits / session) during Stage 2 training, 2) during the first session of Stage 3 training (Stage 3-early), and 3) after mice reached performance criteria during Stage 3 training (Stage 3-late). LFPs were sampled at 2 kHz. Raw traces were visually inspected for noise artifacts, and custom MATLAB functions were used to identify periods of “noise” in the LFP (specifically, “clipping” artifacts, irregularly large amplitude events, and periods of high-amplitude 60 Hz cycles). All traces were then de-trended (removal of low-frequency “drifting” artifacts) with custom MATLAB functions. Multitaper spectral analysis was used for power density estimation and phase coherence via Chronux toolbox routines in MATLAB (http:// chronux.org [37]). For phase-amplitude coupling analysis, phase and amplitude values for frequency pairs were extracted via Morlet wavelet convolution. Phase and amplitude values were then binned (num. bins = 18 per cycle), and mean amplitude values were normalized by dividing each bin value by the summed value over all bins. A modulation index value was then derived by calculating the Kullback-Leibler distance between the observed phase-amplitude distribution and a uniform (null) phase-amplitude distribution[[3]](https://sciwheel.com/work/citation?ids=558928&pre=&suf=&sa=0). For Granger causality analysis, custom MATLAB functions were used to perform multiple lagged autoregressions on LFP traces from the LC and ACC, with the model order for each set of autoregressions determined by the Bayes’ information criteria (BIC). Granger causality values for LC-to-ACC directionality and ACC-to-LC directionality were normalized into a lead index value for ease of comparison between session types and frequency bands.

**Supplementary Figure Legends**

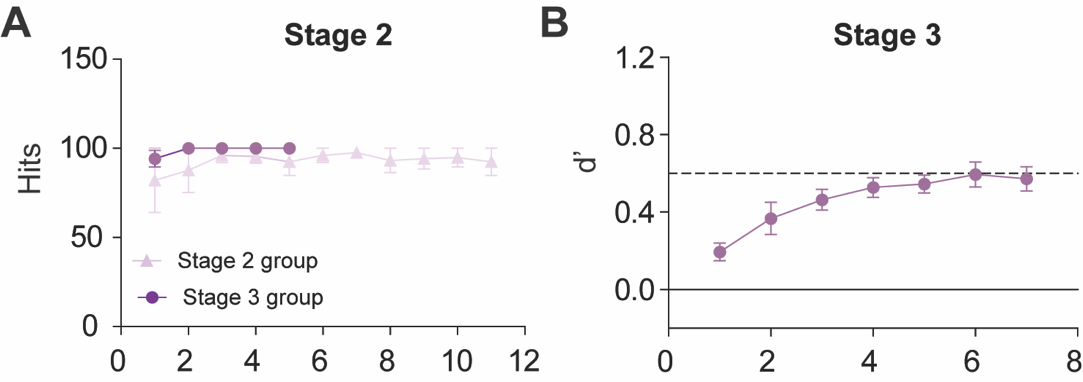

**Supplementary Figure 1. Stage 2 and 3 behavioral performance.** Mice used for c-Fos labeling experiments shown in Figure 1 reached asymptotic behavioral performance during Stage 2 (A) and Stage 3 (B) of rCPT training.

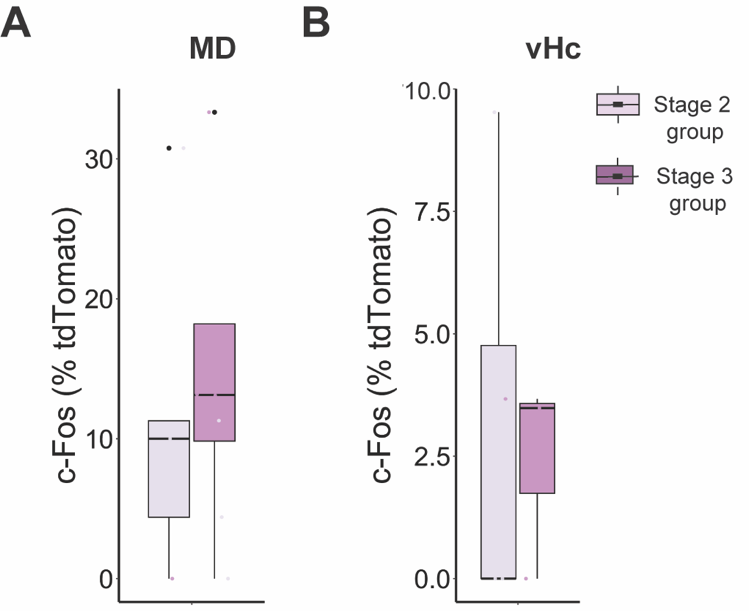

**Supplementary Figure 2. c-Fos expression in projection neurons from mediodorsal thalamus (MD) and ventral hippocampus (vHc) to the ACC.** Boxplots summarizing the quantification c-Fos expression in ACC-projecting (tdTomato+) neurons from the A) mediodorsal thalamus (MD) and B) ventral hippocampus (vHc).

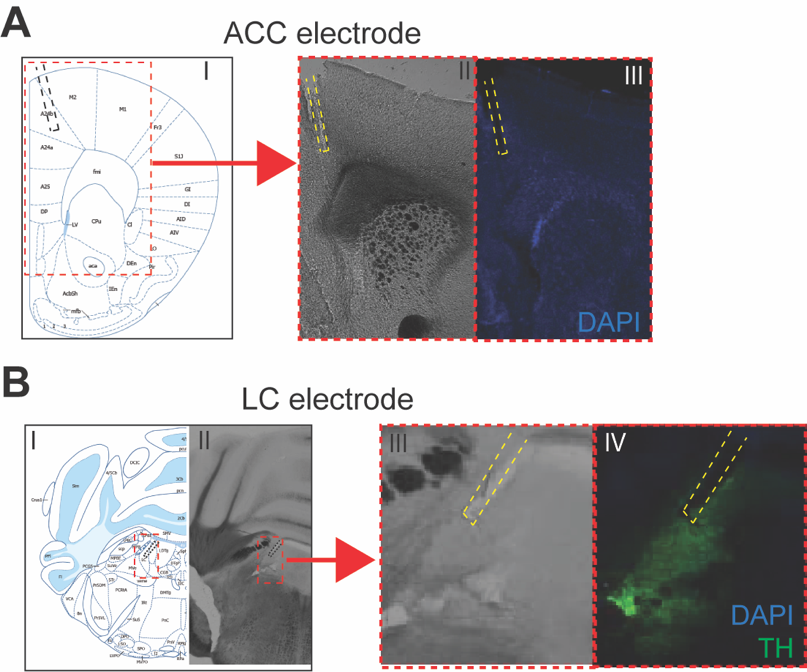

**Supplementary Figure 3. Histological confirmation of electrode placements.**

A) Schematic depicting stereotaxic coordinates for ACC electrode placement (I), and corresponding representative image of bright field (II) and fluorescent staining of DAPI (III) to visualize electrode track within the ACC. B) Schematic depicting stereotaxic coordinates for LC electrode placement (I), with corresponding representative image at low (II) and high (III) magnification bright field pictures of a LC slice, and high magnification fluorescent staining of DAPI (blue) and TH (green) to visualize electrode track within the LC.

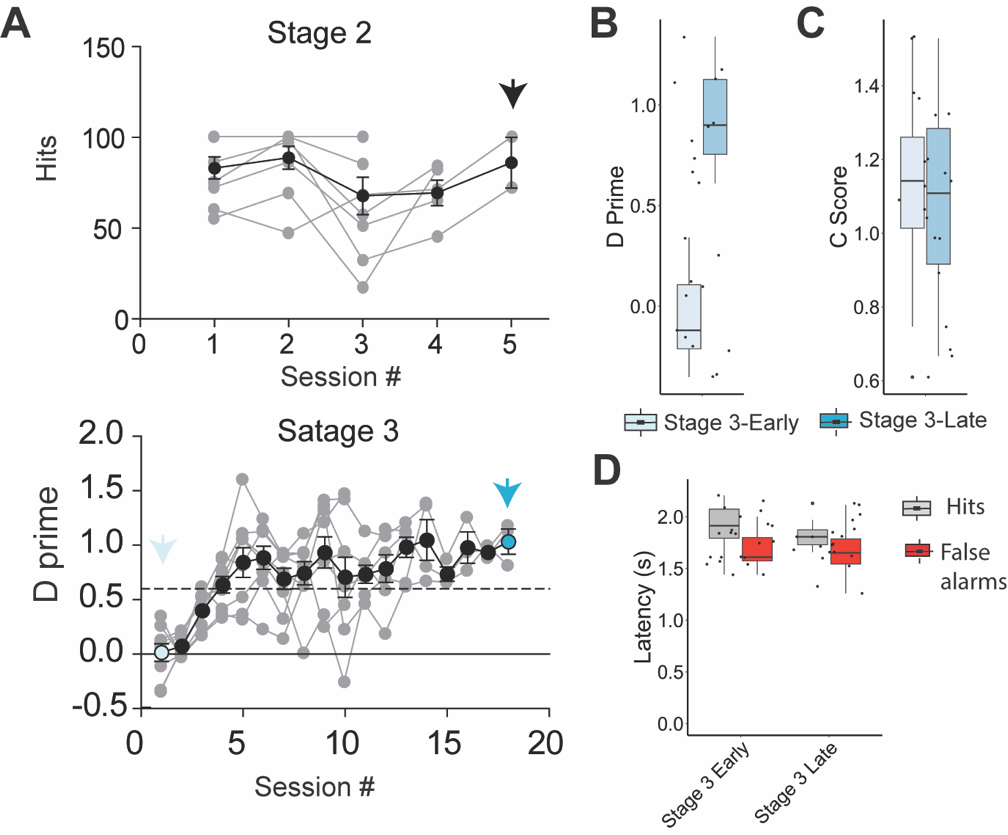

**Supplementary Figure 4. Behavioral performance of mice used for electrophysiological recordings during rCPT training.** A) Time course of behavioral performance during Stage 2 (top) and Stage 3 (bottom) from mice with implanted electrodes that we recorded from. Arrows point performance during recording sessions (black arrow for Stage 2, light blue arrow for Stage 3-early, and dark blue for Stage 3-late recording session). B-C) Boxplots showing discrimination index (d’) and bias response criteria (C score) in Stage 3-early and Stage 3-late. C) Boxplot showing the latency for hits (grey) and false alarms (red) during Stage 3 early and Stage 3-late recording sessions.

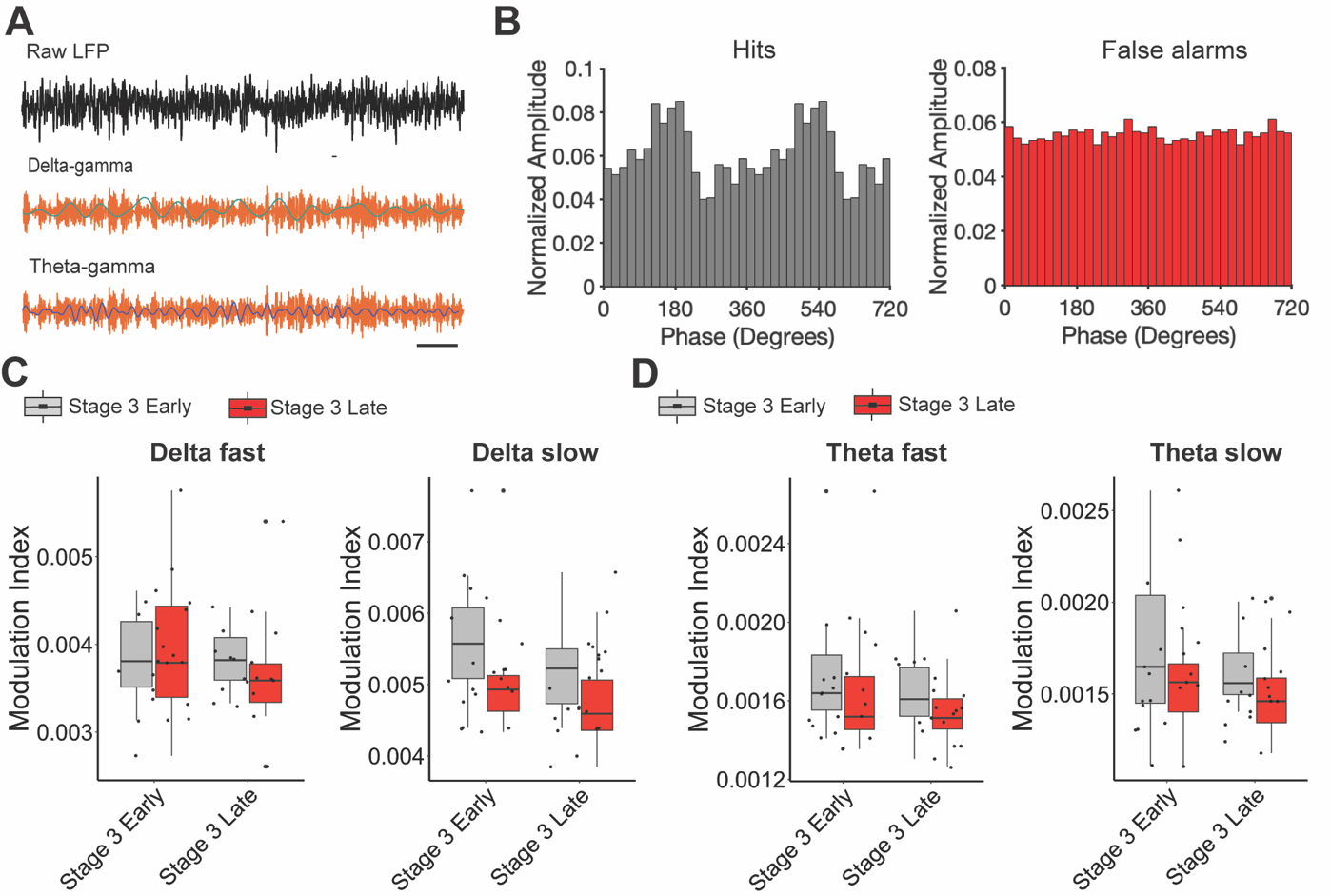

**Supplementary Figure 5.** **Phase-amplitude coupling in the ACC.** A) Representative traces of raw LFP (top), filtered delta (light blue) and gamma (orange; middle), and theta (dark blue) and gamma bands. Scale bar = 0.5 seconds. B) Phase amplitude histograms showing delta-slow gamma coupling of hits and false alarms during Stage 3-early C) box plots showing delta-fast gamma and delta slow-gamma coupling D) Box plots showing delta-fast gamma and delta slow-gamma coupling.
